# Imatinib mesylate induces necroptotic cell death and impairs autophagic flux in human cardiac progenitor cells

**DOI:** 10.1101/2021.04.12.439436

**Authors:** Robert Walmsley, Derek S. Steele, Georgina M. Ellison-Hughes, Andrew J. Smith

## Abstract

The receptor tyrosine kinase inhibitor imatinib mesylate has improved patient cancer survival rates but has been linked to long-term cardiotoxicity. This study investigated the effects of imatinib on cell viability, apoptosis, autophagy and necroptosis in human cardiac progenitor cells in vitro. After 24 hours, imatinib significantly reduced cell viability (75.9±2.7% vs._100.0±0.0%, n=5, p<0.05) at concentrations comparable to peak plasma levels (10 µM). Further investigation showed no increase in caspase 3 or 7 activation. Imatinib also significantly reduced the fluorescence of cells stained with TMRM (74.6±6.5% vs. 100.0±0.0%, n=5, p<0.05), consistent with mitochondrial depolarization. Imatinib increased lysosome and autophagosome content relative to the control, as indicated by changes in acridine orange fluorescence (46.0±5.4% vs. 9.0±3.0, n=7, p<0.001) and expression of LAMP2 (2.4±0.3 fold, n=3, p<0.05) after 24 hours treatment. Although imatinib increased the expression of proteins associated with autophagy, it also impaired the autophagic flux, as demonstrated by the proximity ligation assay staining for LAMP2 (lysosome marker) and LC3II (autophagosome marker), with control cells showing 11.3±2.1 puncta per cell and 48 hours of imatinib treatment reducing the visible puncta to 2.7±0.7 per cell (n=10, p<0.05). Cell viability was partially recovered by autophagosome inhibition by wortmannin, with a 91.8±8.2% (n=5, p>0.05) increase in viability after imatinib and wortmannin co-treatment. Imatinib-induced necroptosis was associated with an 8.5±2.5-fold increase in activation of mixed lineage kinase domain-like pseudokinase. Imatinib-induced toxicity was rescued by RIP1 inhibition relative to the control; 88.6±3.0% vs. 100.0±0.0% (n=4, p>0.05). In summary, imatinib applied to human cardiac progenitor cells depolarizes mitochondria and induces cell death through necroptosis, which can be recovered by inhibition of RIP1, with an additional partial role for autophagy in the cell death pathway. These data provide two possible targets for co-therapies to address imatinib-induced long-term cardiotoxicity.

## Introduction

Receptor tyrosine kinases are proteins critical for many cellular processes and are activated via the transfer of phosphoryl groups, driving the downstream signalling required for cell survival and proliferation. Normally this process is tightly controlled, however when dysregulated it can lead to carcinogenesis. Consequently, over the last two decades these proteins have been extensively investigated, leading to the development of inhibitors to prevent their activation^1^. Tyrosine kinase inhibitors are widely-used anti-oncogenic drugs, with an ever-growing range developed for targeted oncogenic properties, with great success in contributing to event-free patient survival^2^. Imatinib mesylate (IM) is a receptor tyrosine kinase inhibitor (RTKI) currently used to treat patients with chronic myeloid leukaemia and gastrointestinal stromal tumours. It is a relatively selective inhibitor of receptor tyrosine kinases such as c-kit, platelet-derived growth factor, intracellular Abl and chimeric fusion protein BCR-Abl^3,4,5^. The survival of patients presenting with chronic myeloid leukaemia has greatly improved since the introduction of IM treatment^6^.

Although IM has demonstrated great benefits to patients, with a reduced side-effect profile and improved survival, it has been linked to adverse side-effects such as cardiotoxicity, particularly within the aging population^7^. The use of IM may also cause longer-term manifestations of cardiotoxicity, especially within children; such problems have been shown with other chemotherapy drugs such as doxorubicin^8^. As IM is a relatively recent drug discovery, with its first clinical trial in 1998, data on such long-term manifestations have yet to be reported^9^. In an early study, reduced left ventricular ejection fraction (below 50% of baseline) was identified as a cardiotoxic side-effect in patients undergoing IM chemotherapy^10^. However a subsequent study of 1276 cases showed adverse cardiac events after IM treatment was rare: 22 patients suffered congestive heart failure, with age and pre-existing cardiac conditions being contributory factors^11^. Other studies showed an association of IM with hypertrophy and heart failure in patients treated for gastrointestinal stromal tumours, along with elevated levels of natriuretic peptide precursor B^12^. Further evidence of the potential adverse effect of IM was demonstrated in cardiomyocytes in vitro, which exhibited altered calcium transients, cardiomyocyte hypertrophy, mitochondrial dysfunction and cell death^13,14,15^. Our previous study identified damaging effects of IM on adult cardiac fibroblasts, with significant impact on viability and proliferation, alongside changes in expression of growth factors and cytokines (TGF-β1; PDGFD; IL6; IL1β^16^), suggesting that further investigation into the mechanisms of IM-induced cardiotoxicity was required, particularly in the myocyte-supporting cell populations of the myocardial interstitium.

Human endogenous cardiac progenitor cells (hCPCs) have attracted significant interest since their discovery in 2003^17^. One endogenous hCPC population, identified by expression of the receptor tyrosine kinase c-kit (in the absence of the haematopoietic lineage marker CD45), can play a critical role in myocardial tissue maintenance^18,19^. C-kit is a tyrosine kinase receptor expressed on the cell surface: when activated by its ligand stem cell factor, it forms a dimer, initiating the activation of cell proliferation, survival and differentiation pathways^20^. C-kit^+^ hCPCs are capable of self-renewal and development into different myocardial cell types, including vascular smooth muscle and endothelial cells^17–19,21,22^. These cells have also been shown to have a limited ability to form cardiomyocytes. However the extent to which they can do this, particularly in ischaemic tissue, is highly debated ^17,21,23,24^. These cells can however make a valuable contribution to cardiac tissue maintenance via both the replacement of damaged vascular cells and by aiding recovery of injured myocytes through the release of pro-survival growth factors^18,22^. Cardiac progenitor cells have also been shown to be necessary for adult myocardial regeneration in a rodent model of diffuse myocardial injury, involving cardiomyopathy induced by catecholamine excess and removal of proliferating CPCs by the antimitotic agent 5-flurouracil. This combination led to severe heart failure, with functional recovery dependent upon CPCs^18^. Therefore, hCPCs may play a key role in resistance to, or recovery from, RTKI-induced cardiotoxicity by providing protection to cardiomyocytes at risk of RTKI-induced cell death.

To fully understand how RTKIs cause adverse effects on the myocardium, it is important to investigate their impacts on cell death pathways. Apoptosis is a tightly-regulated process defined by the activation of pro-apoptotic factors such as executioner caspases 3 and 7, which eventually lead to DNA fragmentation and cell death^25^. An alternative cell death pathway subject to growing investigation is necroptosis (programmed necrosis), this form of necrosis is regulated by the expression of activated proteins: receptor-interacting serine/threonine-protein kinase 1 (RIP1), receptor-interacting serine/threonine-protein kinase 3 (RIP3) and mixed lineage kinase domain like pseudokinase (MLKL)^26,27,28^. Another relevant cellular process is autophagy, which has been linked to cell death, including the leakage of pro-cell death signals from lysosomes^29^. However, other studies have shown autophagy to be an initiator of other cell death pathways such as apoptosis, albeit not their direct cause^30,31^. Understanding the involvement of autophagy in IM-induced cell death could have significant clinical importance as some studies suggest autophagy as a possible cause for IM resistance in cancer patients^32^. This study addresses the cell death pathways involved in IM-induced toxicity in hCPCs with a view to identifying possible mechanism(s) to overcome IM-induced toxicity.

## Results

### IM reduces cell viability but does not induce apoptosis

To determine whether RTKIs caused cell death within the hCPC population, cells were grown in vitro and exposed to a range of concentrations of IM. Cell viability was decreased by 25% relative to control (n=5, p<0.05) after cells were treated with IM for 24 hours (**Figure 1A**). Following this, the mechanism of IM-induced toxicity was investigated to determine whether cell death was occurring through apoptosis or an alternative pathway. ToPro-3 staining was apparent within the nuclei of hCPCs after treatment with 10 µM and 100 µM IM (**Figure 1B-D**). There was a dose-dependent increase in ToPro-3-positive cell numbers after IM treatment, with a significant increase of 13.6±1.5% after 100 µM IM (n=4, p<0.05) (**Figure 1E**). Neither control cells nor IM-treated cells showed increased caspase 3/7 activity (**Figure 1C)**. Real time RT-qPCR was used to detect changes in gene expression after exposure to 10 µM IM for 24 hours. There were no significant changes in expression of either apoptotic (BAX, caspase 8, PARP, calpain and TNF-α) or necroptosis-associated genes (RIP1, RIP3 and MLKL). The largest expression fluctuations seen were in caspase 8 (1.6±0.5 fold), TNF-α (1.6±0.2 fold) and RIP3 (1.4±0.1 fold) (n=3) **Figure 1F**).

**Figure 1.**
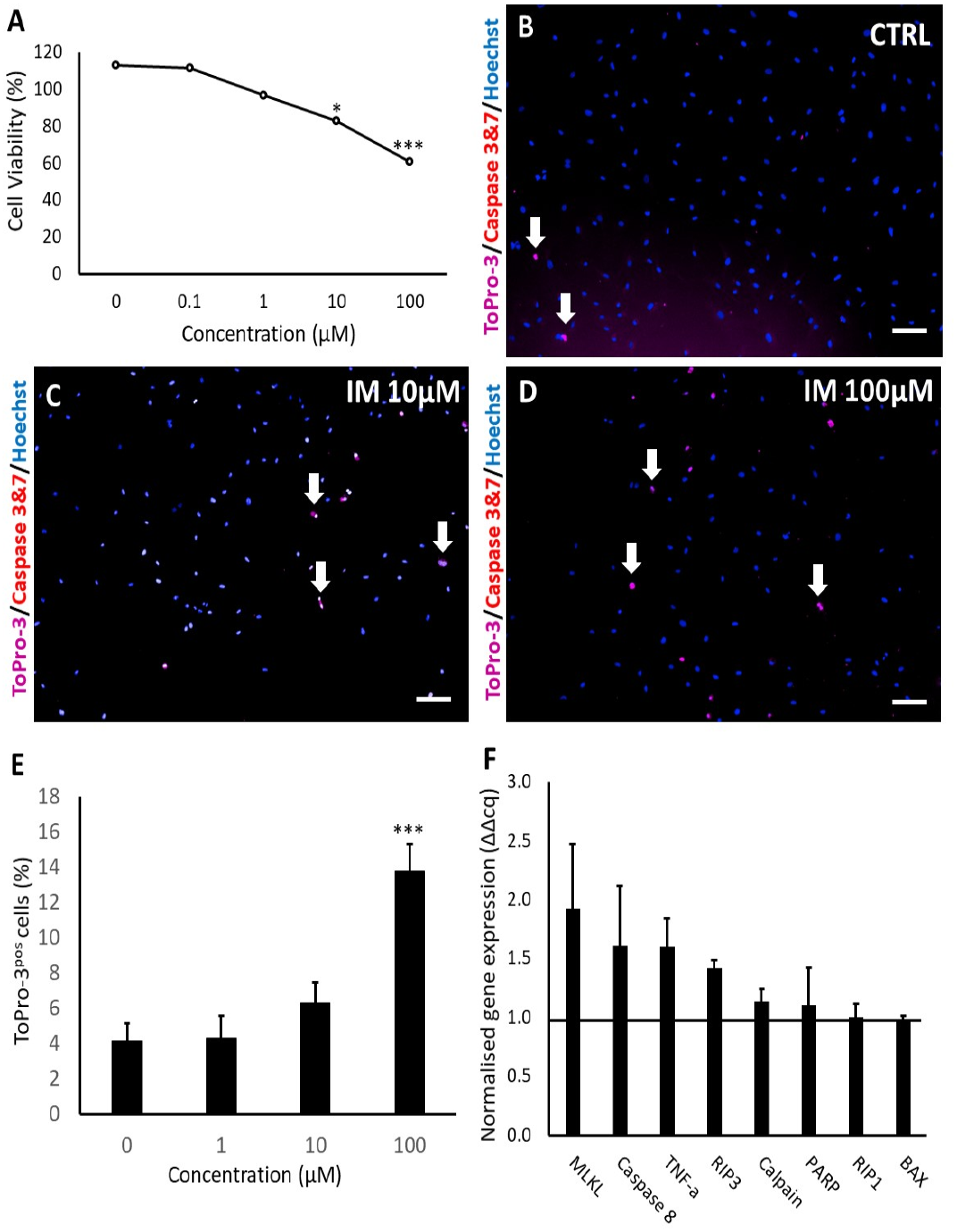
IM reduces hCPC viability but does not induce apoptosis. **(A)** Cell viability was measured by FDA assay, IM treatment ranged from 0.1-100 µM, n=5. **(B-D)** Representative images of live cell staining with ToPro-3 and caspase 3&7 substrate, with increased concentration of IM showing no caspase activity. **(E)** Quantification of ToPro-3 shows an increase in ToPro-3 staining with increased IM concentration, n=4. **(F)** Real time RT-qPCR shows no change in apoptosis-associated gene expression, control=1 (represented by dashed line). Data are mean±SEM, n=3, ***p<0.05 vs. control. Scale bar=100 µm

### IM impairs mitochondrial membrane potential in hCPCs

There was a reduction in the number of cells with high intensity for TMRM after 10 µM IM treatment, with 20% of these cells having a reduced intensity; 20 µM of IM reduced the number of cells with high intensity by 38% and 50 µM IM reduced the number of such cells by 60% (**Figure 2A-D**). Finally the positive control FCCP decouples the mitochondria, reducing by 96% cells within the gated region relative to untreated control cells (**Figure 2E**). These data are reinforced by the changes in mean fluorescent intensity (MFI) of TMRM, with a reduction in average fluorescence of 74.6±6.5% after 10 µM IM treatment and of 49.2±6.8% after 50 µM IM treatment, relative to control (n=6, p<0.05) (**Figure 2F**).

**Figure 2.**
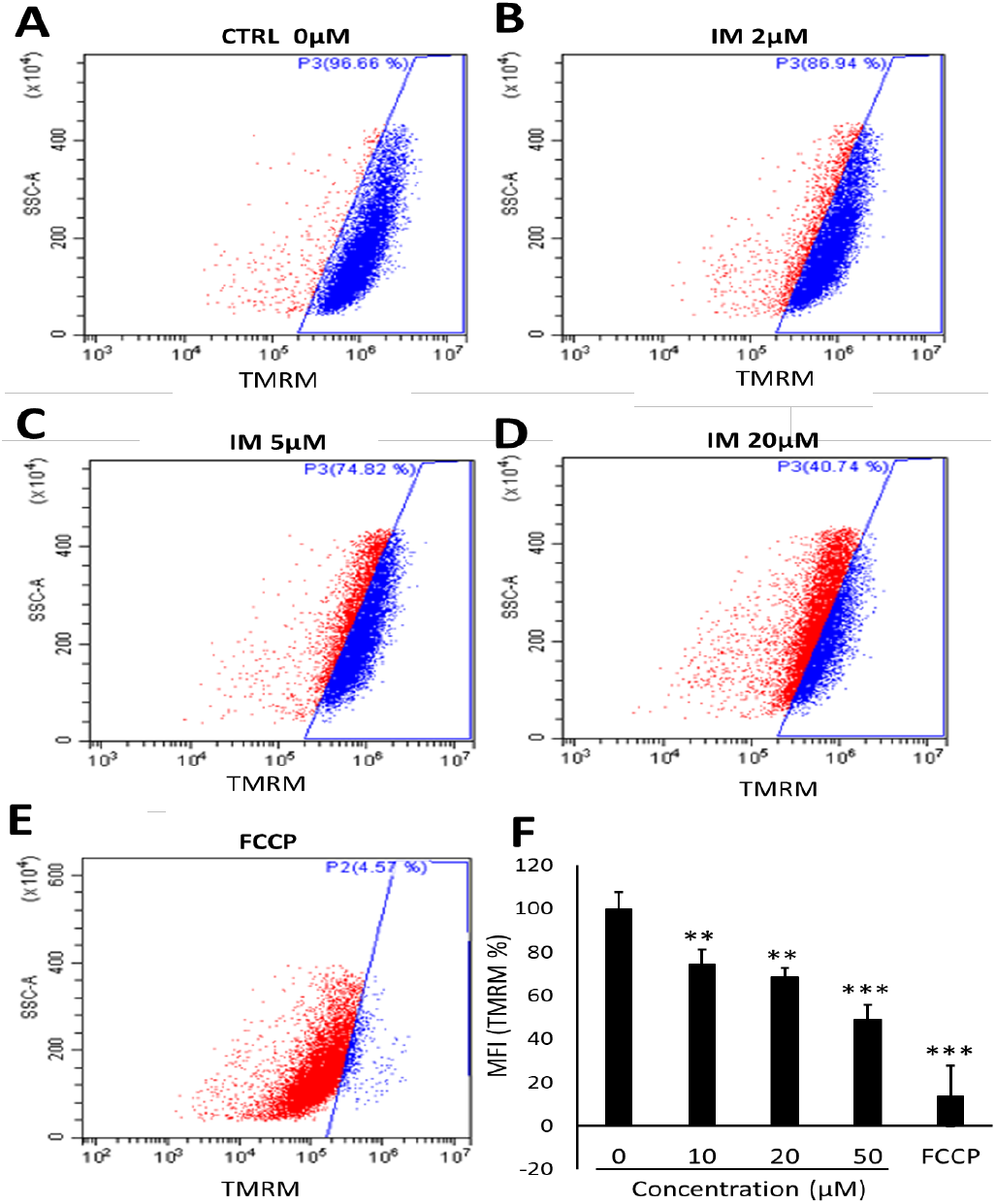
IM reduces hCPC mitochondrial membrane potential. Changes in TMRM fluorescent intensity with increased concentration of IM: **(A)** Control (blue), **(B)** 10 µM IM with slight left shift in intensity (red), **(C)** 20 µM IM **(D)** 50 µM IM and **(E)** positive control 10 µM FCCP **(F)** Quantification of mean florescence intensity (MFI) of TMRM normalised against the control (100%). Data are mean±SEM, n=5, **p<0.01, ***p<0.001 vs. control.

### IM impairs autophagic flux in hCPCs

The cells were exposed to 10 µM IM over periods of 24, 48 and 72 hours, and showed a significant increase in number of acidic organelles (**Figure 3A-D**). Quantification of acridine orange-positive cells showed control cells to be 9.0±3.0% positive, IM-24 hours 55.5±6.4% positive, IM-48 hours 60.0±5.7% positive and IM-72 hours 73.2±3.1% positive (**Figure 3E**). To examine underlying mechanisms, Western blotting was used to analyse expression of LAMP2, a specific lysosomal protein marker. An increase in LAMP2 protein expression was seen in Western blotting, quantified through densitometry, with a 2.4±0.3-fold increase in LAMP2 protein expression after exposure of cells to 10 µM IM for 24 hours (**Figure 3F**).

**Figure 3.**
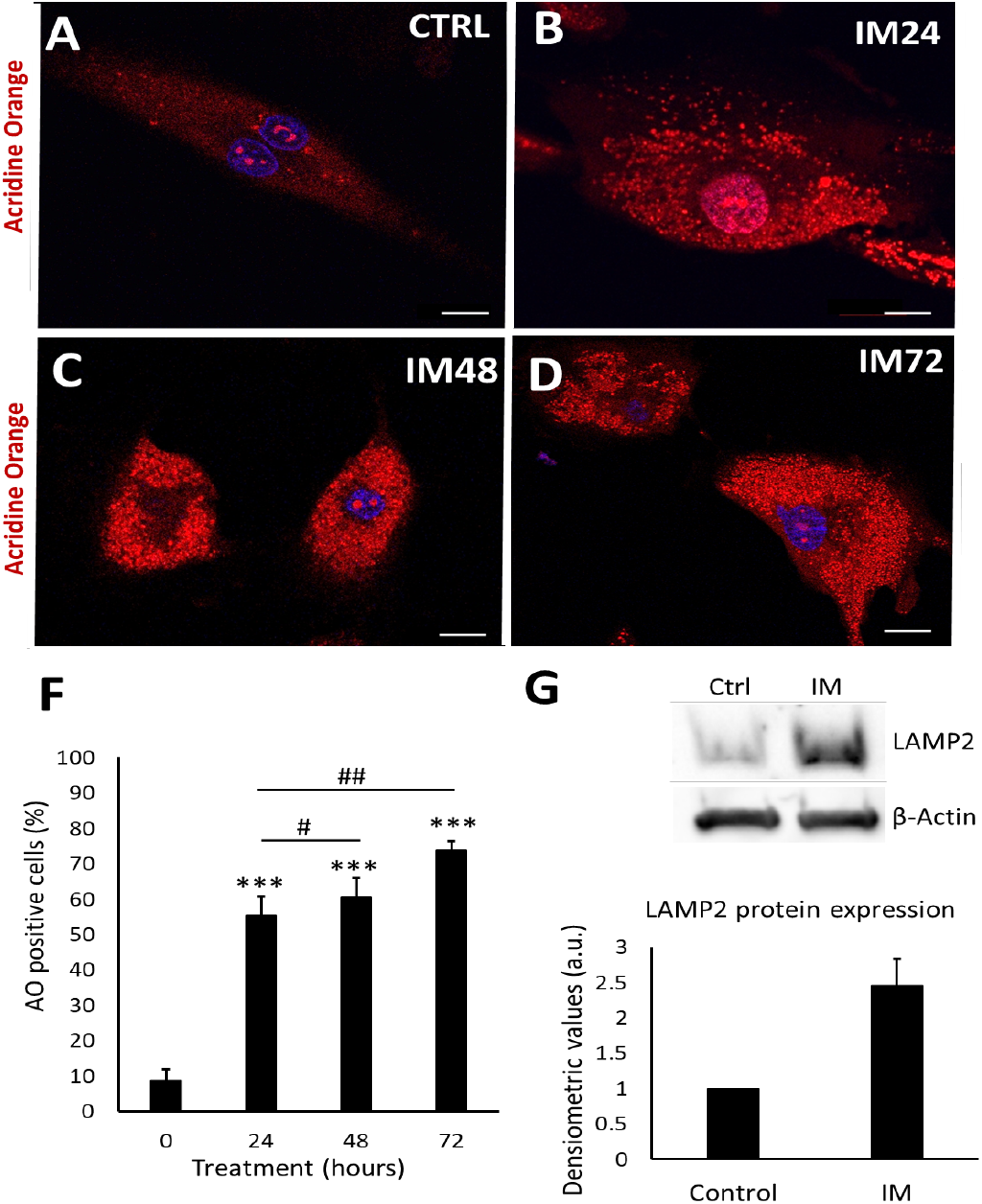
IM induces accumulation of acidic vesicle organelles indicative of autophagy and LAMP2 expression. **(A-D)** Representative images of hCPCs stained with acridine orange, orange staining represents acidic organelles (arrows): control cells, IM (10 µM) 24, 48 and 72 hour treatments. **(E)** Quantification of acridine orange staining in hCPCs after IM treatment (10 µM) relative to control. Data are mean±SEM for percentage of acridine orange-positive cells, n=7. **(F)** Western blot image for LAMP2, control cells and IM-treatment for 24 hours, beta-actin represents loading control. Densitometry for LAMP2 Western blots average signal intensity normalised against control. Data are mean fold change±SEM, n=3, *p<0.05, ***p<0.001 vs. control, #p<0.05, ##p<0.01 vs. IM 24 hours. Scale bar=20 µm.

Having identified that IM increased the density of acidic organelles and lysosome formation, it was important to evaluate the autophagic flux. Autophagic flux can be defined as how well autophagosomes can fuse with lysosomes, in order to remove unwanted proteins such as p62. To measure autophagic flux, Western blotting was used to analyse the lipidation of the autophagosome marker LC3I to LC3II and the removal of P62 protein. A lysosomal inhibitor known as bafilomycin A1 (bafilomycin) was used to prevent protein degradation, therefore if the cell had a functional autophagic flux, there would be an increase in protein levels after bafilomycin inhibition. Western blot analysis of 10 µM IM treatment for 24 hours caused 2.7±1.3 and 1.9±0.0 fold increases in LCII and P62 respectively (p<0.05), which further increased with bafilomycin to 3.5±0.3 and 3.9±0.9 fold increases (P<0.05). However, 10 µM IM treatment for 48 hours caused 2.6±0.2 and 2.8±0.3 fold changes of LC3II and P62, with no significant increases following bafilomycin co-treatment: 2.7±0.1 and 2.6±0.0 relative to the untreated control cells (1.0±0.0) (**Figure 4A-B**).

**Figure 4.**
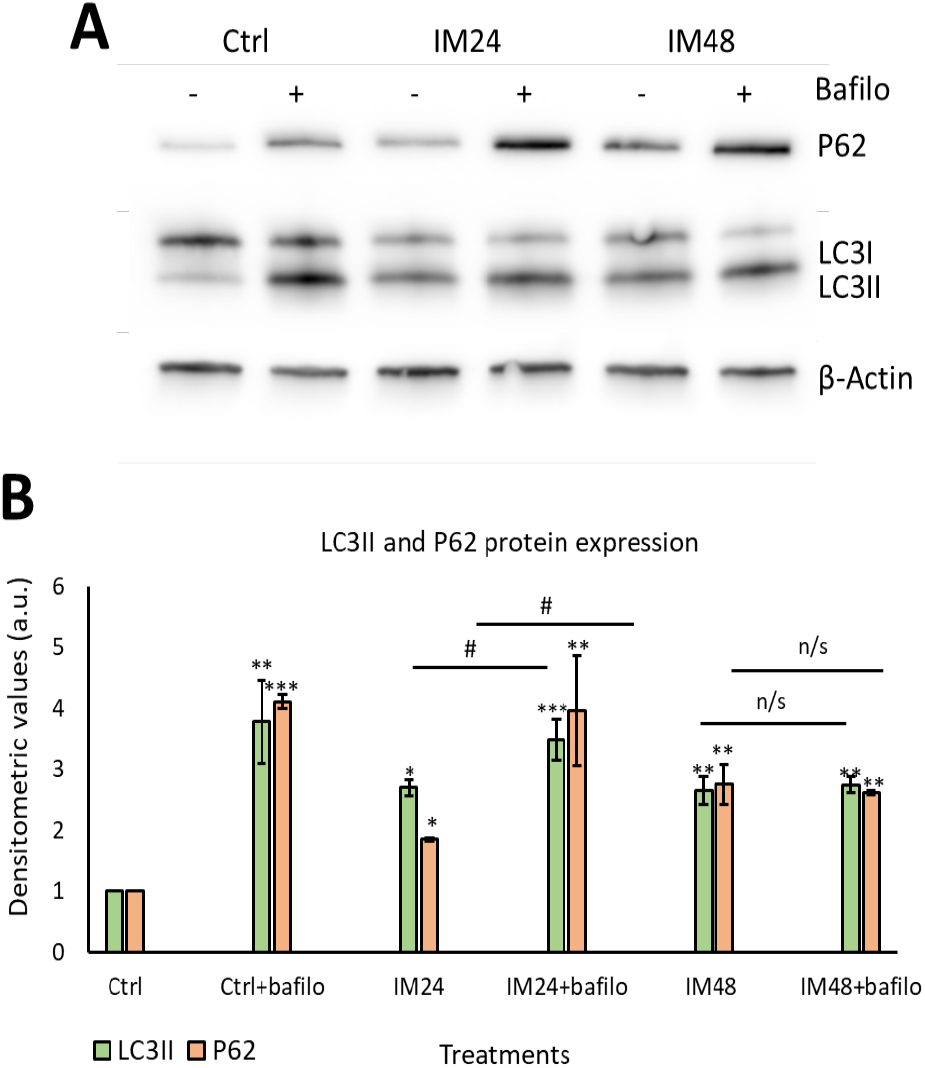
IM impaires the autophagic flux in hCPCs. **(A)** Representative image of Western blotting for P62 and LC3II after IM treatment with/without bafilomycin. **(B)** Quantification of Western blotting for LC3II (green) and P62 (orange). Data are mean±SEM, n=5, *p<0.05, **p<0.05, ***p<0.001 vs. control, #p<0.05 vs. IM 24 hour treatment.

These data were reaffirmed using the PLA assay: this enables a resolution of 50 nm, giving more detailed images to confirm the colocalization between LC3II and p62. Cells were treated with 10 µM IM for 24 or 48 hours, then fixed and stained for LC3II and LAMP2. Cells treated with rapamycin for 4 hours provided a positive control for autophagy. A clear increase in red puncta density can be seen when examining control cells and rapamycin; more puncta can also be seen when comparing control to 10 µM IM treatment for 24 hours. However, there was a clear reduction in red puncta when comparing control cells with those treated by 10 µM IM for 48 hours (**Figure 5A-D**). These appearances were confirmed by quantification of PLA puncta per cell: control cells showed 11.3±2.1 puncta per cell, which increased to 32.1±3.7 after 10 µM IM treatment for 24 hours (p<0.05). The puncta were significantly decreased in number after 10 µM IM treatment for 48 hours: 2.7±0.7 compared to levels after 24 hours (**Figure 5E**) (n=10, p<0.05).

**Figure 5.**
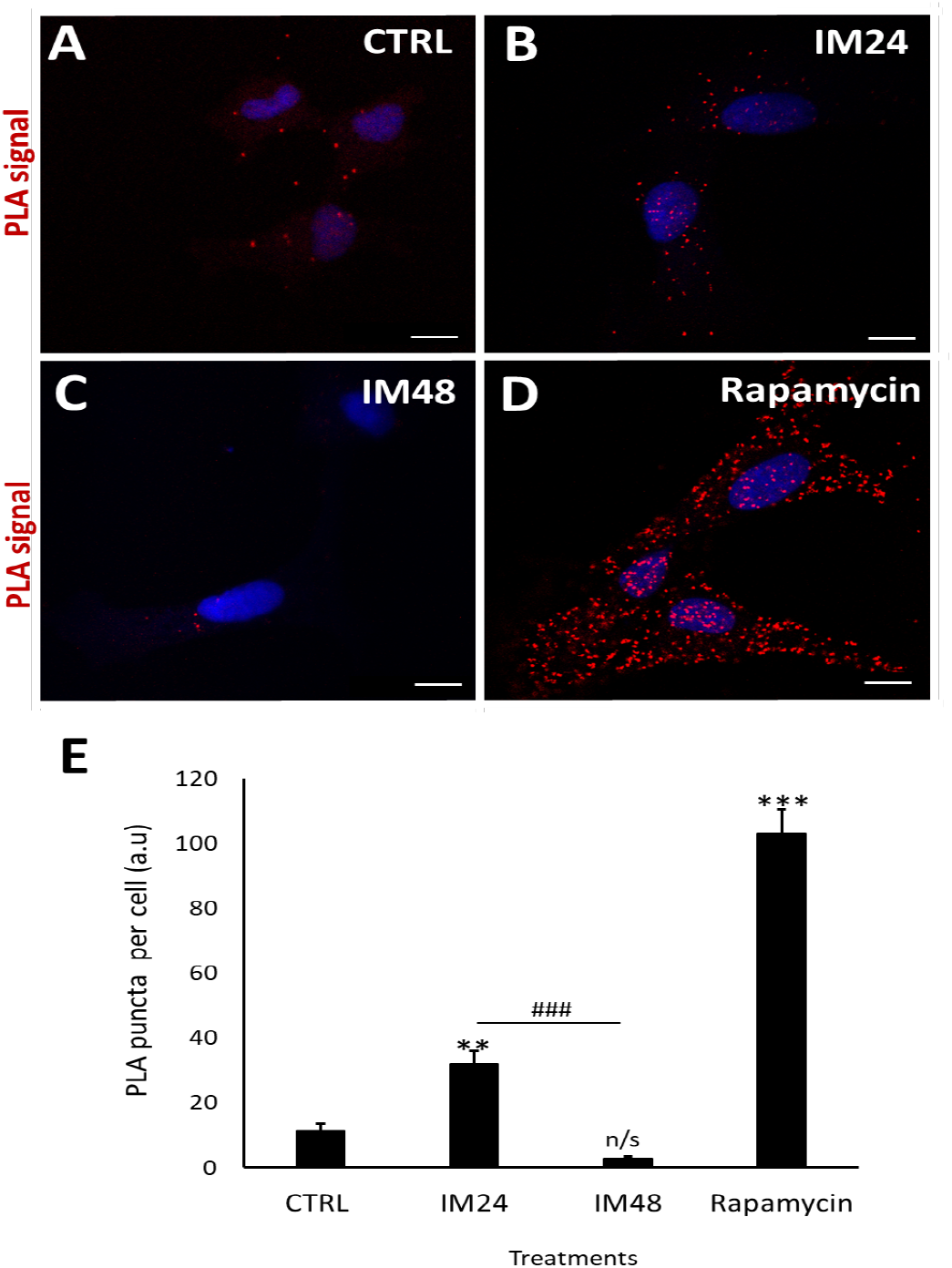
IM reduces LC3/p62 colocalization in hCPCs. **(A-D)** PLA representative images of control cells (low PLA signal), IM 24 hours increased signal, IM 48 hours reduced signal and rapamycin high PLA red puncta (arrows). **(E)** Quantification of PLA puncta after treatment with: IM 24 hours, IM 48 hours and rapamycin 4 hours. Data are mean±SEM, n=10, **p<0.05, ***p<0.001 vs. control, ###p<0.001 vs. IM 24 hour treatment. Scale bar=10 µm.

### Autophagic impairment is not the main contributor to IM-induced hCPC death

Having determined that IM impairs autophagic flux, it was important to identify whether this contributes to cell death. To examine this an autophagosome inhibitor was used (wortmannin): this prevents the initial formation of autophagosomes and is therefore an upstream autophagy inhibitor. Western blotting was carried out to ensure that wortmannin was preventing autophagosome formation, by analysing LC3II expression (**Figure 6A**). Densitometry analysis of LC3II levels identified a decrease in LC3II expression after co-treatment with 10 µM IM and 200 nM wortmannin for 24 hours: 1.5 vs. 2.2 fold when cells were treated with IM only. There was no change when cells were treated with IM for 48 hours and co-treatment with wortmannin: 2.5 vs. 2.3 fold IM only (**Figure 6B**). Following this, cell viability was measured using the FDA assay on treated and control cells: cells treated with IM for 24 hours suffered a 84±1.5% reduction in viability, whereas cells co-treated with IM and wortmannin for 24 hours showed a reduced viability of 91±3.7% (n=3, p<0.05), relative to untreated control values of 100% (**Figure 6C**).

**Figure 6.**
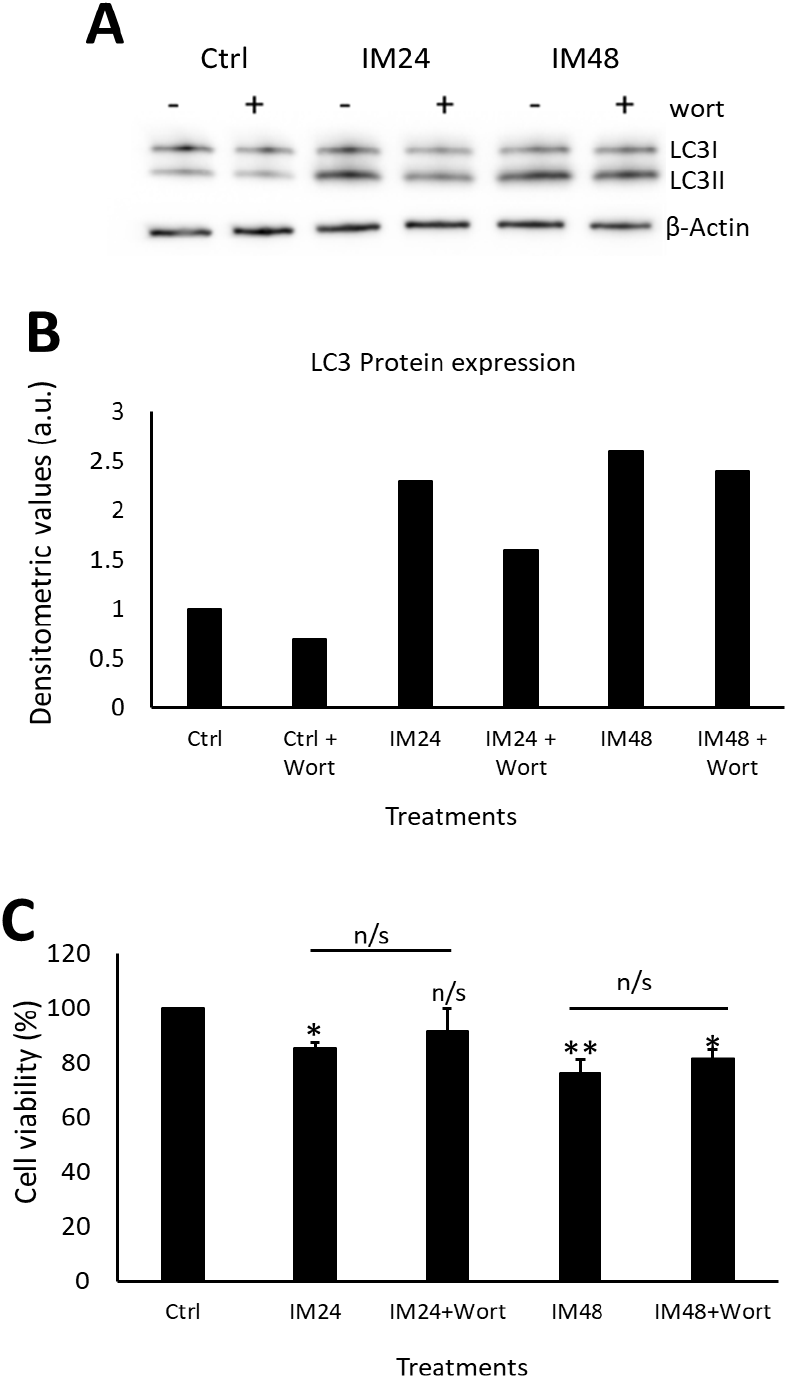
Autophagic flux only partially contributes to IM toxicity in hCPCs. **(A)** Western blot image and densitometric values show LC3II expression after IM treatment for either 24 or 48 hours, with/without wortmannin (wort) co-treatment, n=1. **(B)** CPCs viability after treatment with or without IM and wortmannin treatment for 24 hours. Data are mean±SEM, n=3, *p<0.05, **p<0.01 vs. control, #p<0.05 vs. IM24 or IM48, n/s=not significant.

### IM induces necroptotic cell death within hCPCs, reversed by RIP1 inhibition

Western blotting revealed a clear increase in total MLKL and phosphorylated MLKL after IM treatment (**Figure 7A**). Densitometry of phosphorylated MLKL confirmed this, with an 8.5±2.5 fold increase after 10 µM IM treatment compared to control (**Figure 7B**). Necrostain-1 (nec-1) was used to inhibit the upstream necroptosis initiator RIP1; following nec-1 inhibition cell viability was measured. Compared to control, 10 µM IM treatment for 24 hours reduced cell viability by 78.2±2.5% (p<0.05), whereas 10 µM IM treatment for 24 hours co-treated with nec-1 reduced cell viability by 88.7±3.1% (n=3, p>0.05) (**Figure 7C**).

**Figure 7.**
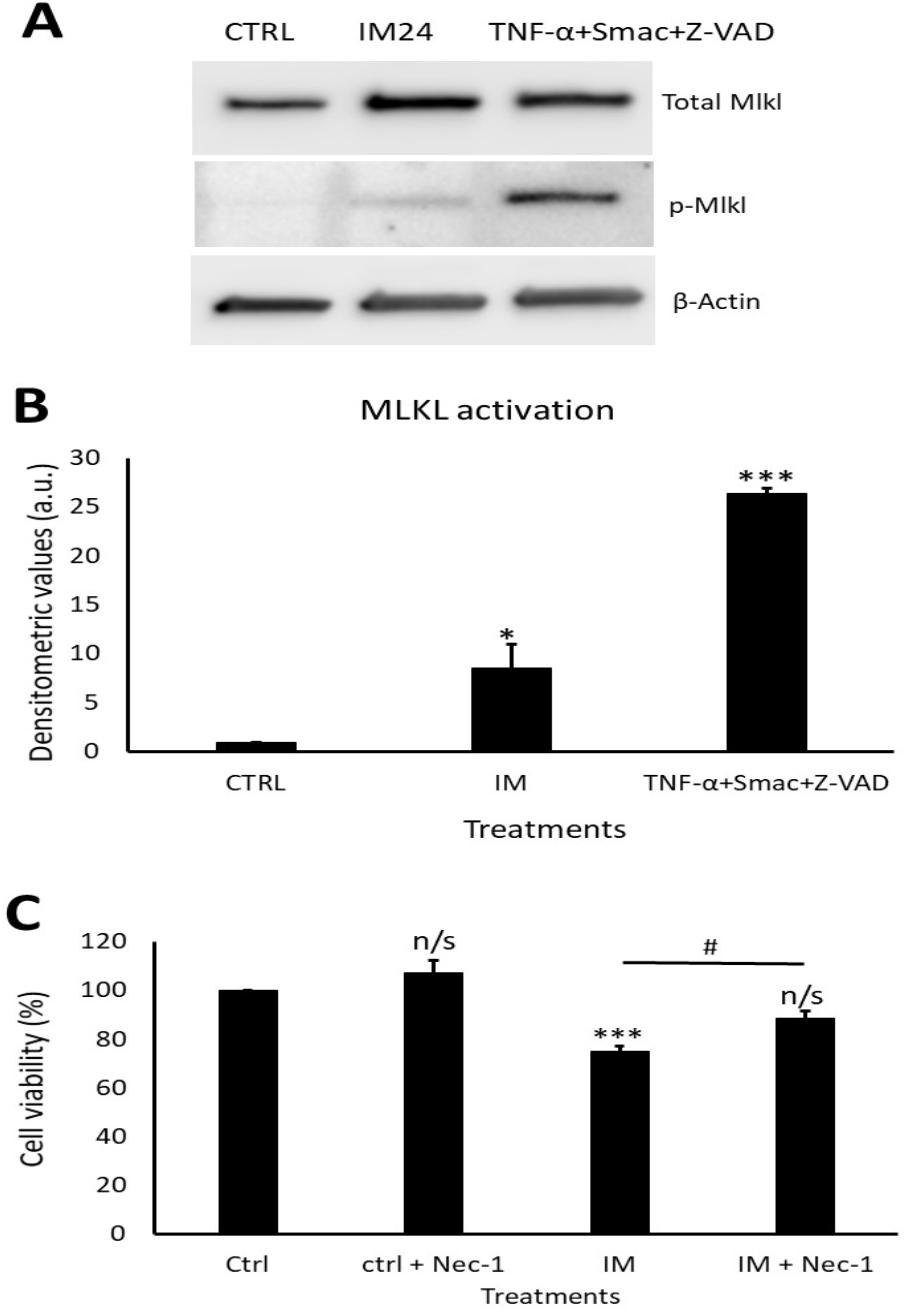
IM induces necroptosis cell death and is rescued by RIP1 inhibition. **(A)** Western blot representative image showing expression of total MLKL and p-MLKL after IM 24 hours treatment and positive control. Densitometric values show MLKL activation after IM treatment and in the positive control, n=3. **(B)** Cell viability analysis of necrostatin (nec-1) inhibition with control cells treated with nec-1, IM alone and IM co-treated with nec-1. Data are mean±SEM, n=4, *p<0.05, ***p<0.001, n/s=not significant vs. control, #p<0.05 vs. IM 24 hour treatment.

## Discussion

This study demonstrates that IM causes significant impacts on the hCPC population. Only a few previous studies have fully investigated the cell death pathways induced by IM, with a very small number addressing IM effects on hCPCs ^16,34–36^. Here we found that IM reduced viable hCPC numbers, but without inducing apoptosis. Previous reports have shown IM induced an increase in caspase 3 and 7 expression, BAX, cytochrome C and TUNEL-positive marking, in cardiomyocytes and cancer cell lines^10,35^. This present study did not find evidence of activated caspase 3 and 7, or increased expression of apoptosis-associated genes. However, there was an increase in ToPro-3 staining, consistent with severe plasma membrane damage and cell death. Further investigation of IM effects on mitochondrial membrane potential were in keeping with previous findings in cardiomyocytes and chronic myeloid leukaemia cells, with IM causing a reduction in membrane potential^10,37,38^. One reason for this could be leakage of lysosomal content such as zinc during autophagy, as previously shown by Li et.al.^39^. Therefore, it was important to analyse the contribution of autophagy in IM-induced cell death.

The number of acidic organelles increased after IM application, which was accompanied by increased expression of the lysosomal structural protein LAMP2. This is similar to findings in previous studies in the mouse neuroblastoma cell line N2a, with one study suggesting IM-induced autophagy is due to inhibition of c-Abl rather than of c-kit or PDGF^40^. There are no studies investigating autophagic flux in hCPCs. Here, autophagic flux was shown to be impaired, so although IM increased the amount of autophagosomes and lysosomes, their ability to fuse was likely impaired. However, when IM-induced autophagy was blocked by the upstream autophagy inhibitor wortmannin, there was no significantly decrease in cell viability compared to untreated cells, suggesting no toxic effect on the CPCs. Wortmannin could therefore be a potential therapeutic target to aid in protection of hCPCs, although the existing data are contradictory. One previous study showed that inhibiting upstream autophagy using 3-methyladenine (3MA) or Atg5 inhibition impaired the chemotherapeutic effect of IM in a range of cancer cell lines^41^. In apparent contradiction, other reports using K562R cells have shown 3MA improved IM-induced cancer cell death^42^. Therefore, further work is required to understand fully if wortmannin could be used to protect hCPCs, while also avoiding inhibition of IM’s chemotherapeutic action.

Due to the impairment of autophagic flux and an increase in p62 protein, this study investigated the involvement of necroptotic cell death in IM-induced toxicity. A previous study by Goodall et.al. showed the presence of p62 protein to be vital for MLKL phosphorylation; if p62 was removed the cell death pathway would switch to an apoptotic cell death^43^. This current study has shown the absence of key apoptotic markers after IM treatment: the expression of phosphorylated MLKL was therefore measured to determine whether IM was inducing cell death through necroptosis. We found an increase in MLKL activation compared to untreated cells. This cell death pathway has yet to be identified within IM-induced cardiotoxicity or within previous cancer cell research. These previous studies have shown IM-induced toxicity to occur through apoptotic cell death^10,35,44,45^, although one study was unable to identify apoptosis in cardiomyocytes^13^. As the viability of hCPCs was rescued using the RIP1 inhibitor nec-1, this could provide a valuable therapeutic target to protect hCPCs from damage and thereby support cardiomyocyte survival. Further experiments would be needed to ensure nec-1 is not involved in the chemotherapeutic action of IM, although previous cancer studies show cell death through apoptosis and not necroptosis; this could however be due to a lack of investigation into necroptosis mechanisms.

In summary, this study demonstrated a range of impacts caused by the clinically-used cardiotoxic drug IM on human endogenous cardiac progenitor cells. The study identified two possible targets to overcome IM-induced toxicity on hCPCs, through either the inhibition of upstream autophagy or via the inhibition of necroptosis (**Figure 8**).

**Figure 8.**
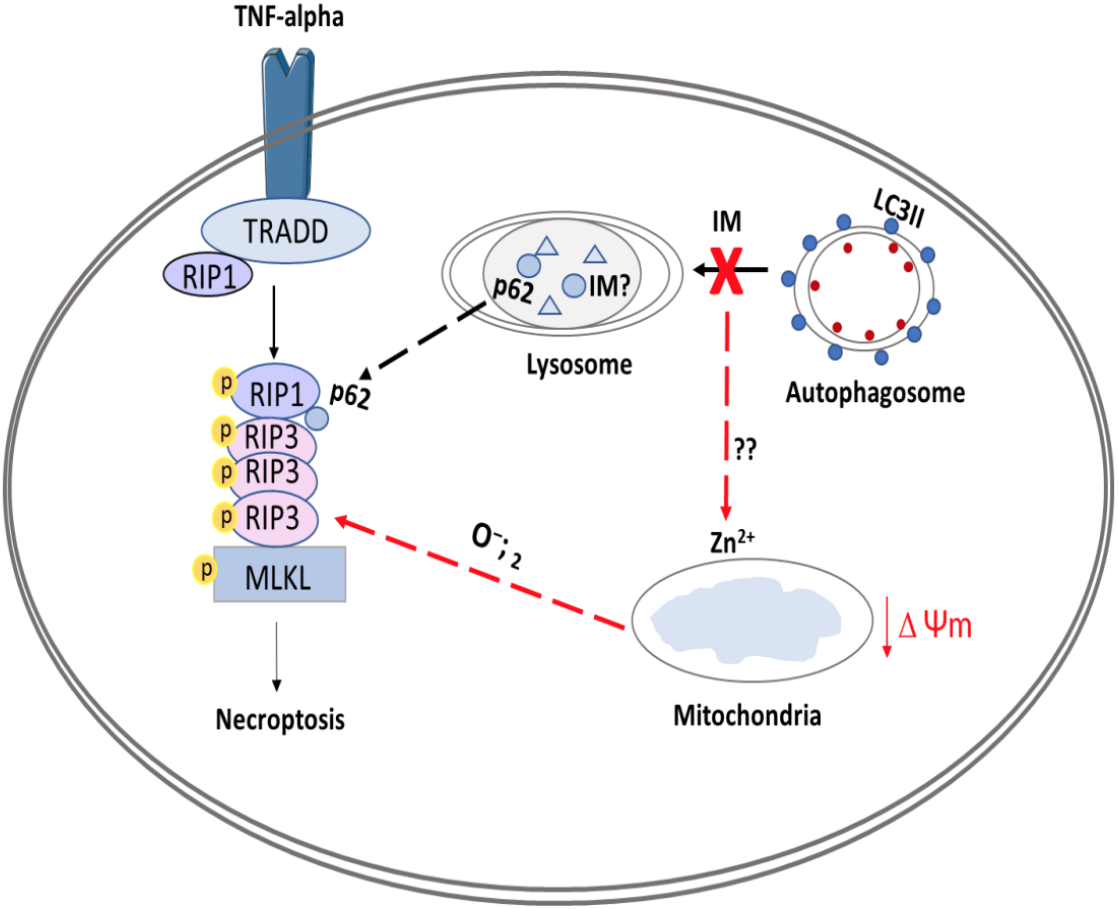
Schematic overview of proposed model for IM-induced cell death in CPCs. The figure shows IM being sequestered by the lysosomes, causing an impaired autophagic flux. This abnormal flux causes the release of zinc ions which affect the ΔΨm leading to production of superoxides (O- ;2);these superoxide ions can help form the ripoptosome. The ripoptosome formation is also aided by leakage of p62 proteins from lysosomes and autophagosomes. Finally, the ripoptosome formation causes the activation/phosphorylation of MLKL and initiates necroptosis cell death.

## Materials and Methods

### Cell culture

All cell culture techniques were performed under aseptic conditions. Growth medium was composed using two solutions: solution A comprised DMEM-F12-Ham’s containing insulin-transferrin-selenium (1% vol/vol), basic-FGF (10 ng/ml), EGF (20 ng/ml) and human leukaemia inhibitory factor (10 ng/ml). Solution B comprised Neurobasal medium supplemented with Glutamax (2% vol/vol), B27 supplement (2% vol/vol) and N2 supplement (1% vol/vol). To prepare growth medium: 45% of solution A and 45% of solution B were combined and supplemented by: embryonic stem cell–qualified FBS (10% vol/vol); penicillin-streptomycin (1% vol/vol); Fungizone (0.1% vol/vol); gentamicin (0.1% vol/vol), then sterilised through a 0.22 µm pore filter. ‘Medium’ is defined as the above, unless otherwise stated.

### Human cardiac progenitor cell isolation

Adult hCPCs were isolated from myocardial tissue obtained during the routine course of cardiac surgery, donated by: Mr Prakash Punjabi (Hammersmith Hospital); Mr. David O’Regan and Mr. Sotiris Papaspyros (Leeds General Infirmary). Briefly, biopsies were weighed and dissected into 0.5-1 mm^3^ pieces and subjected to collagenase type II digestion 0.3 mg/ml (Lorne laboratories) at 37°C: first for 5 min, followed by 7-9 repeat digestions for 3 min, with each post-digest suspension passed through a 100 µm filter. After tissue sample digestion, all collected cells were strained through a 40 µm filter to isolate the smaller cell population. The collected cell fraction was spun at 400 g for 10 min, prior to elimination of debris through Optiprep solution (Sigma). Cells were pelleted by centrifugation through a dual-layered Optiprep suspension: lower layer of 36% (16 ml) and upper layer of 16% (16 ml) at 800 g for 20 min. The cells were then negatively selected for CD45 and positively selected for CD117 (c-kit) using magnetic bead separation (Miltenyi Biotec). After selection, the hCPCs were re-suspended in medium and plated onto a coated 6-well plate for conditioning. Cells were cultured in a humidified tissue culture incubator at 37°C, 5% CO_2_, 1% O_2_ and passaged upon reaching 70-80% confluency. All reagents were sterilized through a 0.22 µm filter.

### Imatinib treatment

Imatinib (Synkinase) was either added at the peak plasma level equivalent concentration of 10 µM (for reverse transcription qPCR, acridine orange staining, proximity ligation assay and Western blotting experiments), or in a dose-dependent manner of 1 µM-100 µM depending on experimental conditions (cell viability, mitochondrial membrane potential staining, live cell staining for high content image analysis).

### Cell viability

The fluorescein diacetate (FDA) cell viability assay was performed as previously described^33^. Briefly, 5×10^3^ hCPCs were plated into each well of a 96-well plate and treated with IM in medium for 24 hours. After treatment, medium was removed and replaced with 5 µg/ml FDA (in 1% ethanol, 9% PBS and 90% DMEM-F12), incubated for 10 min at 37°C. Fluorescent readings were taken using a Varioskan Flash plate-reader (v.4.00.53, Thermo Fisher) at excitation/emission wavelengths of 485/520 nm. Cell viabilities of treatment groups were calculated as a percentage of the untreated control group. For cell viability with autophagy inhibitor, cells were left to attach for 24 hours before 10 µM IM treatment and co-treatment with 200 nM wortmannin for 24 and 48 hour time points before applying FDA as described.

### Live cell staining

Live cell staining for high content image analysis: cells were added to a CellCarrier-96 Ultra Microplates (Perkin Elmer), 1000 cells/well. Fam/Cas dye (1:150) was added and incubated for 1 hour and washed with apoptosis wash buffer (Life Technologies V35118; diluted 1:10). Hoechst 1:200, ToPro-3 1:1000 and TMRM 1:100,000 (Invitrogen) added and incubated for 10 min, then replaced with medium (without indicators) and analysed by an Operetta platform (Perkin Elmer). Results were interpreted using the Columbus™ software (Perkin Elmer).

### Mitochondrial membrane potential staining

In order to evaluate the effect of IM on mitochondrial membrane potential, flow cytometry was used to analyse the retention of a cell permanent cationic dye called tetramethylthodamine methyl ester perchlorate (TMRM). Cells were seeded onto 100 mm tissue culture dishes until 80% confluent; TMRM (1:300,000) was added for 10 min and the cells were then washed with PBS before being detaching using Accutase (Sigma). Following this, the cells were centrifuged at 300 g for 5 min and then re-suspended in a PBS wash; the cells were then pelleted at 300 g for 5 min. Finally, cells were re-suspended in incubation buffer: PBS (Ca^2+^ and Mg^2+^ free, Invitrogen 14190-136); 2.5 g bovine serum albumin (Sigma A9418); 1% (v/v) Penicillin/Streptomycin (Invitrogen 15140-122); 0.1% (v/v) Fungizone (Invitrogen 15290-018); 0.1% (v/v) Gentamicin (Sigma G1397) and analyzed using a Cytoflex S flow cytometer (Beckmans) using 561/585 nm filter. Data were interpreted using Cytexpert and Excel. Mitochondrial membrane potential was calculated using the fluorescence intensity of TMRM of treated cells relative to untreated.

### Acidic organelle staining

To determine the role of autophagy in IM-induced cell death, acridine orange (ImmunoChemistry Technology) was used to determine if the drug increased acidic properties within the cell (acidic organelles), indicating the presence of increased lysosome content. Cells were plated into a 12-well plate and allowed to reach confluency. Following this, 5 µM acridine orange was added to medium for 30 min. Cells were then washed once with PBS and imaged using confocal microscopy (Zeiss LSM880). The number of positive cells were counted using Image J (National Institutes of Health).

### Reverse transcription quantitative PCR (RT-qPCR)

Cells were plated onto 100 mm dishes and treatments added at a clinically comparable concentration. Cells were pelleted and re-suspended in 1 ml of PBS with antibiotics (1% (v/v) Penicillin/Streptomycin (Invitrogen 15140-122); 0.1% (v/v) Fungizone (Invitrogen 15290-018); 0.1% (v/v) Gentamicin (Sigma G1397)) and centrifuged 1500 rpm for 5 min. Briefly, RNA was then isolated using the QIAshredder and RNeasy mini kit (Qiagen), according to the manufacturer’s guidelines. The RNA sample purity was tested using a Nanodrop 2000 (Thermo Fisher). The iScript cDNA synthesis kit and BioRad CFX96 Real Time system were used to generate 100 µl cDNA from 2000 ng RNA. mRNA sequences for all primers were obtained from the NIBC nucleotide database and assessed using Primer-BLAST software (NCBI). All PCR primers (Sigma) were reconstituted in molecular grade water (Sigma) to make 100 µM long-term stocks and 5 µM working stocks. SYBR green (BioRad), molecular grade water (BioRad) and forward and reverse primers were added to 96-well plate, with 2 µl cDNA per well added to give a total volume of 20 µl per well. Samples were run for 40 thermal cycles with real-time imaging of PCR product generation (CFX96, BioRad). Data were analysed using CTX manager software (BioRad) and Microsoft Excel, with data representing ΔΔCq values.

### Proximity ligation assay

The proximity ligation assay (PLA), gave visualisation of two proteins within 40 nm of each other (this indicates higher probability of direct binding). The Duolink kit was used (Sigma DUO92002, DUO92004, DUO92008), with cells cultured on 8-well glass chamber slides (Thermo Fisher 155409) and treated with 10 µM IM for 24 or 48 hours, or rapamycin 400 nM for 4 hours. Cells were fixed with 2% paraformaldehyde for 10 min at room temperature, then permeabilised with 0.1% Triton X-100 and blocked with goat serum (Thermo Fisher, 10000C) for 1 hour at room temperature. Cells were then incubated with primary antibodies: 1:200 LAMP2 (Novus) and 1:200 MAP1LC3B Antibody (CSB-Cusabio, cat.no. PA887936LA01HU) overnight 4°C. For secondary incubation, PLA (+) and PLA (-) probes were incubated for 1 hour in a humidified incubator. For each treatment 8 µl of PLA (+) and 8 µl of PLA (-) were added in 24 µl of blocking buffer (total volume of 40 µl) and placed onto treatment wells. The cells were then washed with PLA wash buffer A and incubated with ligation reaction (40 µl) which includes 8 µl of ligation buffer, 1 µl ligase in 31 µl of dH_2_O for 30 min in humidified chamber. The cells were washed 3 times with PLA wash buffer A and incubated with the amplification reaction (8 µl amplification buffer; 0.5 µl DNA polymerase in 31.5 µl of dH_2_O) for 100 min in a humidified incubator at 30°C. Finally, cells were washed three times with PLA wash buffer A and 0.01% wash buffer B for 10 min. The results were visualised using confocal microscopy (Zeiss LSM880). Following image acquisition, PLA puncta staining was manually counted for each condition.

### Western blotting

Cells were plated on 100 mm tissue culture dishes and treated with: 10 µM IM for 24 hours; bafilomycin 400 nM for 3 hours; 10 µM IM for 24 hours and bafilomycin 400 nM for the last 3 hours of treatment; untreated aside from medium change (control). For necroptosis positive control, cells were incubated with 20 ng/ml TNF-α (Peprotech), 100 nM SMAC mimetic Birinapant (BioVison), 20 µM V Z-VAD(OMe)-FMK (Tonbo biosciences) for 8 hours. After treatment, cells were lysed using RIPA buffer (Sigma) and quantified using Pierce™ Rapid Gold BCA Protein Assay Kit (Thermo Scientific). Protein was loaded (10-50 µg per well, depending on protein of interest) and transferred using semi-dry turbo blotter (Bio-Rad). Blots were placed into 5% blocking solution containing primary antibodies and were incubated overnight at 4°C (**Table 2**). The blots were then washed three times (10 min per wash) in TBS-Tween, following this blots were placed in blocking solution containing the relevant secondary antibody, either anti-rabbit (Cell Signaling) or anti-mouse (Cell Signaling), at 1:1000 dilution for 1 hour at room temperature. Blots were developed using West Pico plus (Thermo Fisher) and imaged on the automated developer G-box (Syngene). Densitometry was analyzed using Image J and Excel, with treatments normalized against control groups.

**Table 1.**
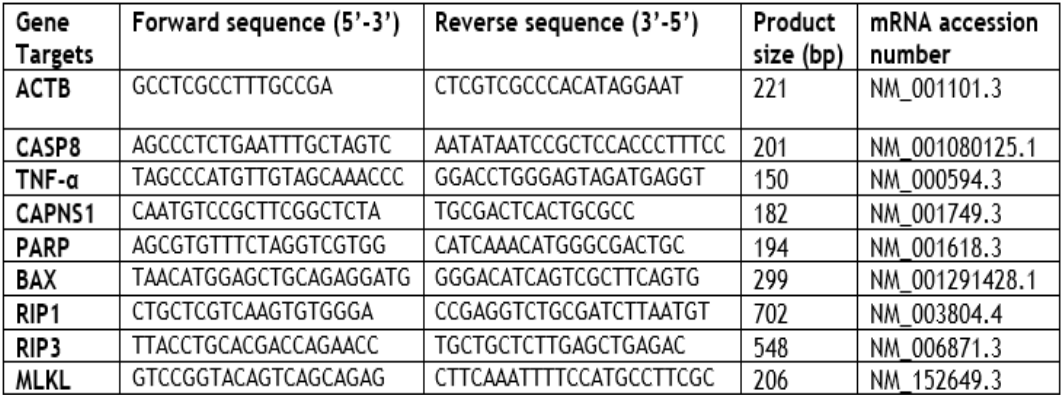
Table of primer pairs for real time RT-qPCR. Forward and reverse primer sequences, product size and mRNA accession number for each gene target.

**Table 2.** Table of primary antibodies. List of antibodies used for Western blotting and ICC staining (including dilutions used and host species).

| Host | Target | Company | Dilution |
| --- | --- | --- | --- |
| Rabbit | Anti-human LC3 | Novus | 1:4000 |
| Rabbit | Anti-human MAP1LC3B | Cusabio | 1:200 |
| Mouse | Anti-human LAMP2 | Novus | 1:1000 |
| Rabbit | Anti-human p-MLKL | Abcam | 1:1000 |
| Rabbit | Anti-human total MLKL | Genetex | 1:1000 |
| Rabbit | Anti-human beta-actin | Cell signalling | 1:2000 |
| Rabbit | Anti-human p62 | ProSci Inc. | 1:1000 |

### Statistical analysis

One-way ANOVA was used for comparison of mean data between groups for analyses of cell viability, live cell staining, mitochondrial membrane potential staining and PLA assay, followed by post hoc Tukey test for comparisons between groups. Two-tailed, unpaired Student’s t-test on ΔΔCq values was employed to compare target gene expression between each treatment and the respective control values.

## Author Contributions and Notes

The authors declare no conflict of interest.

## Acknowledgments

We acknowledge the assistance of the Flow Cytometry and Imaging Facility, University of Leeds. Adult human myocardial samples were provided by: Mr Prakash Punjabi (Hammersmith Hospital); Mr. David O’Regan and Mr. Sotiris Papaspyros (Leeds General Infirmary). This work was carried out with funding from the Leeds Anniversary Research Scholarship award and the School of Biomedical Sciences, University of Leeds. We are also grateful for the assistance of Ms. Miriam Hurley (University of Leeds), Mr. Tim Munsey (University of Leeds), and the cellular cardiology group (University of Leed

## Notes

### Competing Interest Statement

The authors have declared no competing interest.

